# EMT Induction In Normal Breast Epithelial Cells By *COX2*-Expressing Fibroblasts

**DOI:** 10.1101/2024.09.26.615069

**Authors:** Minwoo Kang, Somayadineshraj Devarasou, Tae Yoon Kwon, Jennifer H. Shin

## Abstract

**Background:** The tumor microenvironment (TME) plays a pivotal role in cancer progression, with cancer-associated fibroblasts (CAFs) significantly influencing tumor behavior. Especially, elevated *COX2* expressing fibroblasts within the TME, notably in collagen-dense tumors like breast cancer, has been recently emphasized in the literature. However, the specific effect of *COX2*-expressing CAFs (*COX2*^+^ CAFs) on neighboring cells and their consequent role in cancer progression is not fully elucidated.

**Methods:** We induced *COX2^+^* fibroblasts by forcing the fibroblasts forming aggregates to undergo Nemosis as a proxy for *COX2*^+^ CAFs. This approach enabled us to simulate the paracrine interactions between *COX2*^+^ CAFs and normal breast epithelial cells via conditioned media from *COX2^+^* fibroblasts. We developed an innovative *in vitro* platform that combines cell mechanics-based analysis and biomolecular assays to study the interactions between *COX2*^+^ fibroblasts and normal breast epithelial cells. By focusing on the mechanical characteristics of the cells and the EMT marker expressions, we aimed to elucidate the paracrine mechanisms through which *COX2*^+^ CAFs influence the tumor microenvironment.

**Results:** Our *in vitro* findings reveal that *COX2*^+^ fibroblasts, through conditioned media, induce significant changes in the mechanical behavior of normal breast epithelial cells, facilitating their transition towards mesenchymal types. This transition was corroborated by increased expression of mesenchymal markers. By drawing parallels between *COX2*^+^ fibroblasts and *COX2*^+^ CAFs, we established a positive feedback loop involving *COX2^+^* CAFs, prostaglandin E2 (PGE2), and the EP4-*SNAI1* axes.

**Conclusion:** This study advances our understanding of the potential mechanisms by which *COX2^+^* CAFs influence tumor progression within the breast tumor microenvironment (TME) through controlled *in vitro* investigations. By integrating cell mechanics-based analysis, biomolecular assays, and innovative *in vitro* cell-based modeling of *COX2^+^* CAFs, we have delineated the contributory role of these cells in a controlled setting. These insights lay a groundwork for future studies that could explore the implications of these findings *in vivo*, potentially guiding targeted therapeutic strategies.

## Introduction

The tumor microenvironment (TME) is a complex ecosystem of various cells, including immune cells, endothelial cells, and most notably, fibroblasts, all of which play a crucial role in shaping the evolutionary path of tumors through continuous interaction and signaling with cancer cells^1–3^. Among these, fibroblasts, particularly in their activated form known as cancer-associated fibroblasts (CAFs), are emerging as key contributors to tumor progression. CAFs have a well-documented role in promoting cancer progression through both physical and molecular interactions. Studies, such as those by Zhang et al. ^4^, highlight their involvement in processes like aggregate formation and epithelial-mesenchymal transition (EMT). By orchestrating changes in the extracellular matrix (ECM) and secreting growth factors, CAFs support cancer cell survival and metastasis during tumor development.

The epithelial-mesenchymal transition (EMT) is a critical process in cancer metastasis and progression, wherein epithelial cells acquire mesenchymal characteristics, resulting in a loss of epithelial traits and an increase in migratory capabilities. Epithelial cells, characterized by specific cell junctions such as adherens junctions, gap junctions, and apical-basal polarity, undergo significant changes during EMT, losing these features and adopting back-front polarity indicative of enhanced migratory function^5^. While EMT is widely recognized in contexts such as embryonic morphogenesis and wound healing, it is particularly well-studied in cancer biology^6^. As cancer advances, epithelial cancer cells begin to exhibit mesenchymal traits, becoming more invasive through EMT activation^7–10^. However, this transition is not a binary switch between fully epithelial and fully mesenchymal states^6^. Cells often express a combination of both epithelial and mesenchymal markers, making it challenging to fully capture this phenotypic spectrum through biomarkers alone. To gain a more comprehensive understanding of EMT, it is essential to consider additional factors, such as the biophysical characteristics of the cells, which may provide deeper insights into the complexity and heterogeneity of this process.

Prostaglandin-endoperoxide synthase 2 (*PTGS2*), commonly referred to as cyclooxygenase-2 (*COX2*), is the gene responsible for prostaglandin biosynthesis^11^. Two isoforms of cyclooxygenase exist in humans: *COX1*, which is constitutively expressed in most tissues and regulates the conversion of arachidonic acid to prostaglandins for routine physiological functions^12^, and *COX2*, which is induced during inflammatory responses and is typically absent in normal cells. *COX2* is widely recognized as a biomarker for various cancers^13,14,15^, with elevated levels serving as a distinguishing indicator of tumor presence. Overexpression of *COX2* in cancer cells not only promotes carcinogenesis but also correlates with poorer patient survival^16–20^, contributing to increased resistance to chemotherapy and radiotherapy^21,22^.

Recent research has highlighted the presence of high *COX2* expression within the tumor microenvironment (TME), particularly in collagen-dense tumors and cancers such as colorectal, intestinal, and nasopharyngeal cancers^23–26^. Additionally, a significant association has been found between *COX2*-expressing breast tumors and the number of cancer-associated fibroblasts (CAFs)^27^. In colorectal cancer, CAFs have been identified as the main sources of both *COX2* and prostaglandin E2 (PGE2)^28,29^. Despite this, the impact of elevated stromal *COX2* expression or *COX2*-expressing CAFs (*COX2*^+^ CAFs) on neighboring cells and tumor progression remains insufficiently explored.

PGE2, the end product of *COX2* activity, is a key mediator of inflammation and tissue regeneration. Like *COX2*, persistently elevated levels of PGE2 are strongly linked to various human malignancies. Numerous studies suggest that PGE2 plays a role in promoting cancer cell growth, proliferation, survival, angiogenesis, and metastasis^30–33^, making it a potential prognostic biomarker for cancer^34–36^. PGE2 exerts its effects through four E-series prostaglandin receptors (EP 1-4), engaging in autocrine and paracrine signaling, with responses varying across different cell and tissue types. For instance, while PGE2 inhibits epithelial-mesenchymal transition (EMT) in Madin-Darby canine kidney (MDCK) cells, it promotes EMT in nasopharyngeal cancer cells^25,37,38^.

Nemosis is a process in which fibroblasts become activated and form clusters under certain in vitro conditions^38^. For example, fibroblasts have been shown to form spheroids in response to Bowes melanoma line-conditioned medium. However, there is currently no direct evidence that nemosis occurs at tumor sites in vivo^39,40^. Given that fibroblasts form aggregates at melanoma sites through paracrine interactions with melanoma cells, it is plausible that other tumors could similarly induce a nemosis-like process in nearby fibroblasts via paracrine signaling^41–43^. Nemosis is characterized by increased production of inflammatory mediators, proteinases, and growth factors, making nemotic fibroblasts resemble CAFs. High COX2 expression is considered a hallmark of the nemotic state^44^. Based on this, *COX2*^+^ fibroblasts undergoing nemosis were used as a model for *COX2*^+^ CAFs, given their accessibility and the ability to study cellular mechanisms in a controlled environment, while acknowledging the complexity and heterogeneity of CAF behavior in vivo.

To explore the paracrine signaling dynamics between *COX2*^+^ fibroblasts and neighboring normal breast epithelial cells, we simulated these interactions using conditioned media from *COX2*^+^ fibroblasts, also referred to as nemotic fibroblasts (NF). Enzyme-linked immunosorbent assay (ELISA) revealed that the conditioned media from *COX2*^+^ fibroblasts (NF) contained elevated levels of PGE2, the primary product of *COX2*, compared to the negative control. Further analysis demonstrated that the NF-induced conditioned media (Nemo-CM) were capable of initiating EMT in normal breast epithelial cells. A combination of cell mechanics-based approaches and biomolecular assays was used to assess EMT. The findings of this study support the hypothesis that PGE2, derived from *COX2*^+^ fibroblasts in the tumor stroma, can drive the EMT of normal breast epithelial cells, providing strong evidence for a positive feedback mechanism involved in breast cancer progression.

## Materials and Methods

### Cell culture and preparation of conditioned media

MCF10A cells (ATCC No.CRL-10317) were cultured in DMEM/F12(GIBCO) supplemented with 5% horse serum (Invitrogen 16050-122), 500 ng/ml hydrocortisone (Sigma-Aldrich H0888), 100 ng/ml cholera toxin (Sigma-Aldrich C8052), 20 ng/ml epidermal growth factor (EGF, Peprotech), 10ug/ml insulin (Sigma-Aldrich I1882), and 1% penicillin-streptomycin (PS, Invitrogen 15070-063). Human adult dermal fibroblasts (HDFs, Lonza) were cultured in DMEM (WelGene) supplemented with 10% fetal bovine serum (FBS, WelGene), and 1% penicillin-streptomycin (WelGene).

To induce nemosis, 100 μl of cell suspension drops, containing 1 × 10^4^ HDFs, were seeded on Ultra-Low Attachment 96 well plate (Corning). Each well contained one fibroblast spheroid. After four days of culture, the media (Nemo) were collected and filtered before use (0.2 μm Nalgene syringe filter). For negative control media from flat 96 well plate (Control), the same amount of cell suspension drops was seeded, and the media were collected and filtered before use (Fig. 1a).

**Fig. 1.**
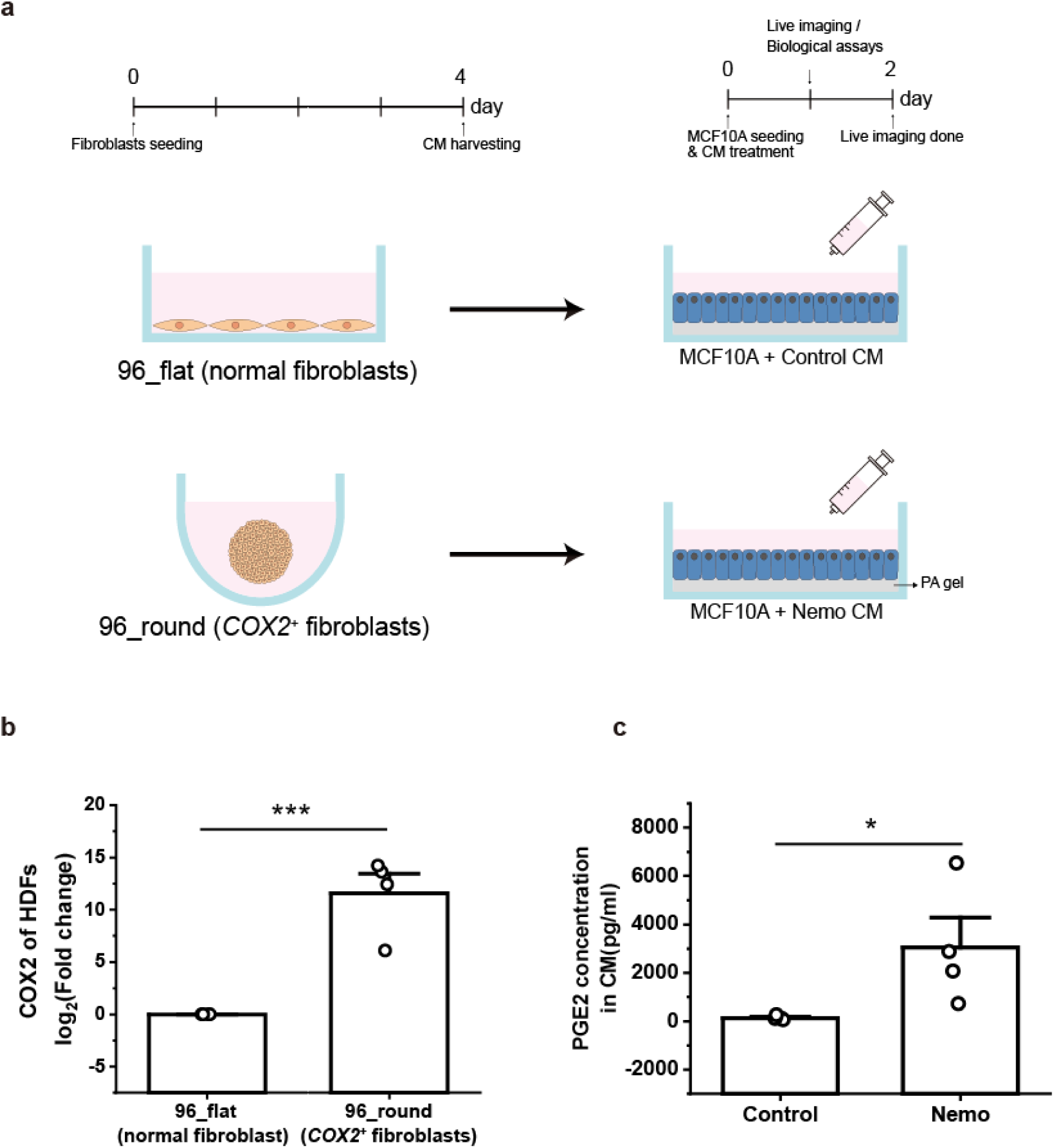
Mimicking paracrine signaling of *COX2*^+^ fibroblasts in vitro. (a) Schematic diagram of experimental design. (b) mRNA expression level of *COX2* of fibroblasts cultured on 96-well flat-bottom (96_flat) and 96-well round-bottom (96_round) cell culture plates. n=4 (c) PGE2 concentrations in conditioned media (CM) harvested from 96-well flat-bottom (Control) and 96-well round-bottom (Nemo) cell culture plates. n=4.

### Enzyme-linked immunosorbent assay

To quantify PGE2 concentrations in conditioned media, conditioned media (control and Nemo) were collected after 4 days of culture. PGE2 concentrations in the conditioned media were measured by an enzyme-linked immunosorbent assay (ELISA) kit (Prostaglandin E2 Parameter Assay Kit, KGB, R&D Systems) according to the manufacturer’s instructions.

### Polyacrylamide gel preparation

Cells were cultured on masked polyacrylamide (PA) gel substrates (Young’s modulus = 3 kPa, thickness = 100 μm) that were prepared following the protocols as previously described ^45^. Briefly, the 24 μl of PA gel drop mixed with red fluorescent beads (diameter = 0.5 μm; FluoSpheres; Life Technologies) was polymerized in a silane-treated glass bottom dish (Spl) under a cover slip (diameter = 18 mm; VWR). The PA gel surface was functionalized with 1 mg/ml of Sulfo-SANPAH (sulfosuccinimidyl-6-(4-azido-2-nitrophenylamino) hexanoate) in warm 50 mM HEPES buffer (Life Technologies) for immobilization of 100 μg/ml collagen type I (PureCol, Advanced BioMatrix).

### Cell monolayer patterning and time-lapse microscopy

Cellular islands were patterned for cell monolayer expansion using polydimethylsiloxane (PDMS; Sylgard 184; Dow Corning) stencil with 700 μm-diameter holes. The PDMS stencil coated with 2% Pluronic F-127 solution (Sigma-Aldrich) at 37 °C for 1 hour. The prepared PDMS stencil was covered on the PA gel substrate. 200 μl aliquot of MCF10A cell-suspension (density = 5×10^5^ cells/ml) was loaded on the pattern. The sample was then incubated at 37 °C / 5% CO_2_ for 30 minutes. After the cell stabilization, cell-suspension was drained to remove the floating cells then conditioned media was treated for 24 hours (control and Nemo) (Fig. 1(a)). All experiments were performed on the Axio Observer.Z1/7 (Carl Zeiss) microscope equipped with a climate-controlled chamber (37 °C and 5% CO_2_). Phase-contrast and fluorescence images were taken every 10 min for 24 hours using a 5x objective lens and 1x optovar magnification changer (Carl Zeiss).

### Fourier transform traction microscopy (FTTM) and Monolayer stress microscopy (MSM)

Tractions of the cellular monolayer were measured using the unconstrained Fourier transform traction microscopy (FTTM) with corrections for substrates of finite thickness, as described in previous reports ^46–48^. To calculate monolayer stress, we used the monolayer stress microscopy (MSM) approach, which is detailed by previously published references ^49,50^. Briefly, intercellular stress was calculated based on the balance of tractions across the cellular monolayer according to Newton’s second law. The principal stresses, *σ_min_* and *σ_min_,* can be converted from the internal stress tensor. For each local point on the cellular monolayer, the local average normal stress *σ_min_* is defined as (*σ_min_*+ *σ_min_*)/2, and it represents the scalar tension within the monolayer.

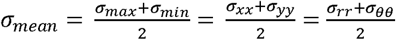

### Single-cell morphological and migratory features quantification

Morphological features of single-cell images were quantified with *imageJ* ^51^ by manually tracking the cell boundary. The aspect ratio, circularity, and solidity were calculated with the following expressions.

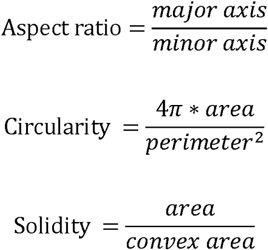

To quantify the migratory features of single-cells, the centroids of single cells were manually tracked with *ImageJ*. After acquiring the centroids of single cells of time series image stacks, mean-squared displacement (MSD), instantaneous velocity, and directional ratio were calculated.

### Immunofluorescence and the EP4 antagonist

Cells were washed with 1X DPBS (Dulbecco’s Phosphate buffered saline, Welgene) and fixed for 15 min in 4%(v/v) paraformaldehyde. After washing three times with PBS, cells were permeabilized by 0.2% (v/v) Triton-X (Sigma-Aldrich) 100 for 10 min and blocked by 3% (v/v) bovine serum albumin (BSA) for 1 h in 4°C. After washing in PBS, cells were incubated with diluted primary antibody (Anti-Vimentin, 1:100, BD Pharmingen™ #550513) overnight, followed by the incubation with secondary antibodies (Alexa Flour 488, 1:100, Invitrogen) for 1 hour in room temperature. Actin filaments were stained with phalloidin (Alexa Fluor 568-phalloidin, 1:100, Invitrogen). For nucleus staining, 4’,6-diamidino-2-phenylindole (DAPI, 1:50,000, Molecular Probe) was treated in the cells for 3 min at room temperature. Cells were then imaged using multichannel fluorescence microscopy (Carl Zeiss).

The EP4 receptor antagonist L-161,982 (SML0690, Sigma-Aldrich) was used for the PGE2 receptor inhibition study. For monolayer expansion experiments, MCF10A cells were first seeded and incubated for 30 minutes with MCF10A culture media for attachment. Then, the attached cells were pre-incubated with serum-free media (DMEM supplemented with 1% PS) with either dimethyl sulfoxide (DMSO) only or L-161,982 (10 μM in DMSO) for preincubation of 1 hour. After the preincubation, cells were washed with PBS and cultured with Nemo-CM for 24 hours.

### Quantitative Reverse Transcription PCR

At the end of the experiments, total RNA was isolated from fibroblasts or MCF10A cells with RNAiso Plus reagent (Takara Bio, Japan) according to the manufacturer’s instructions. Extracted RNAs were reverse transcribed to cDNA using iScript cDNA Synthesis Kits (Bio-Rad, USA) and Biometra T-personal Thermal Cycler for the synthesis. Quantitative Reverse transcription PCR (qRT-PCR) was carried out in duplicates with iQ SYBR green supermix (Bio-Rad, USA) and a Bio-Rad CFX96, real-time detection system. Glyceraldehyde 3-phosphate dehydrogenase (GAPDH) was used for the reference gene. ΔΔC_t_ values were used for hypothesis testing and used to express log_2_(Fold Change) relative to a control group. ΔC_t_ = C_t_(reference gene) – C_t_(gene of interest), ΔΔC_t_ = ΔC_t_(target sample) – ΔC_t_(reference sample). C_t_: Threshold cycle The following primers were used:

GAPDH (For: CTGGGCTACACTGAGCACC, Rev: AAGTGGTCGTTGAGGGCAATG), COX-2 (For: CTGGCGCTCAGCCATACAG, Rev: CGCACTTATACTGGTCAAATCCC), CDH1 (For: CGAGAGCTACACGTTCACGG, Rev: GGGTGTCGAGGGAAAAATAGG), CDH2 (For: TCAGGCGTCTGTAGAGGCTT, Rev: ATGCACATCCTTCGATAAGACTG), VIM (For: AGTCCACTGAGTACCGGAGAC, Rev: CATTTCACGCATCTGGCGTTC), SNAI1 (For: TCGGAAGCCTAACTACAGCGA, Rev: AGATGAGCATTGGCAGCGAG), SNAI2 (For: CGAACTGGACACACATACAGTG, Rev: CTGAGGATCTCTGGTTGTGGT), TWIST (For: GTCCGCAGTCTTACGAGGAG, Rev: GCTTGAGGGTCTGAATCTTGCT), ZEB1 (For: GATGATGAATGCGAGTCAGATGC, Rev: ACAGCAGTGTCTTGTTGTTGT), ZEB2 (For: CAAGAGGCGCAAACAAGCC, Rev: GGTTGGCAATACCGTCATCC).

### Statistical analysis

We assessed the normality of our data using the Shapiro-Wilk test. For hypothesis testing, the Student’s t-test or Mann-Whitney U test was used depending on the normality of the data. P-values < 0.05 were considered as statistically significant. Statistical tests were performed using *jamovi* version 1.6.23.0 (The jamovi project (2021), https://www.jamovi.org)

## Results

### Validation of the simulated *COX2*^+^ fibroblasts through fibroblast aggregates by the comparative expression of *COX2* of fibroblast aggregates and PGE2 concentrations in conditioned media

To evaluate the validity of using *COX2*^+^ fibroblasts formed via in vitro aggregation of fibroblasts as a proxy for *COX2*^+^ CAFs, we first constructed the fibroblast aggregates by culturing the fibroblasts on the low-attachment surface of 96-well round-bottom cell culture plates. Then, *COX2* expression level was checked within fibroblast aggregates and PGE2 concentrations in conditioned media. For the control group, the same number of fibroblasts were cultured on the standard 96-well flat-bottom cell culture plates (Fig. 1a).

To determine *COX2* expression level in fibroblasts, qRT-PCR was performed on the fourth day of culture, which confirmed a significantly higher mRNA expression of *COX2* in aggregated fibroblasts cultured on a low-attachment surface (96_round) compared to those cultured in 2D on a normal surface (96_flat) (Fig. 1b). Next, ELISA analysis confirmed the significantly elevated PGE2 concentrations in the conditioned media (CM) harvested from *COX2*^+^ fibroblasts (Nemo-CM) (Fig. 1c). These results suggest that Nemo-CM can serve as a surrogate for mimicking the paracrine interactions between *COX2*^+^ CAFs and neighboring cells within the TME. To further investigate the impact of this simulated signaling on tumor microenvironment dynamics, we exposed normal breast epithelial MCF10A cells—a model widely used for studying breast cancer genetics^52–54^—to both control and Nemo-CM, and observed their phenotypic changes from cell-mechanical and biomolecular perspectives.

### The collective behavior of cells implies the EMT of normal breast epithelial cells via Nemo-CM

To observe the collective behavior of cells, we established monolayer circles using MCF10A cells. The cells were patterned in circular islands on collagen I-coated polyacrylamide (PA) gel using a PDMS stencil of a circular shape. After cells settled on the hydrogel, the culture medium was changed to CM (control or Nemo). After 24 hours, the stencil was removed to allow cells to settle and form a coherent monolayer (Fig. 2a). When the monolayer came into contact with free space due to stencil removal, the patterned cells began to expand radially. We identified a distinctive behavior of monolayers with different CM conditions (Fig. 2b-g). MCF10A monolayers cultured with Nemo-CM expanded much faster and more aggressively (Supplementary Video 1). We quantified the expansion of monolayers after 16 hours of expansion (Fig. 2b). In addition to the expansion speed, we verified the different cell-cell interactions under various conditions. At the 16-hour mark, snapshots revealed disrupted cell-cell interactions in the Nemo-CM-treated sample. In contrast, the control group maintained robust cell-cell contacts (Fig. 2c), supporting the phenotypic transition of cells when cultured with Nemo-CM. The cells appeared to have undergone EMT based on the increased migratory capacity, the loss of stable cell-cell interactions, as well as the loss of apical-basal polarity in disseminating cells. To further investigate the biophysical characteristics of cellular monolayers, we employed Fourier transform traction microscopy (FTTM) and monolayer stress microscopy (MSM) to analyze tractions and intercellular stresses (Fig. 2d-g). At the initial point of the free expansion, Nemo-CM-treated monolayers showed higher inward traction force (blue-colored) at the edge compared to the control group, indicating that cells at the edge are pulling themselves toward the free space (Fig. 2d). In addition, the Nemo-CM-treated group also had more inward traction regions in the center core, evidenced by more blue-colored area, which suggests that cells at the core also endeavored to migrate outward like the expanding front. This trend was maintained throughout the experiment. This result supports our speculation on EMT and aligns with the previous report that TGF-β-induced EMT enhances the traction forces of A549 cells^55^.

**Fig. 2.**
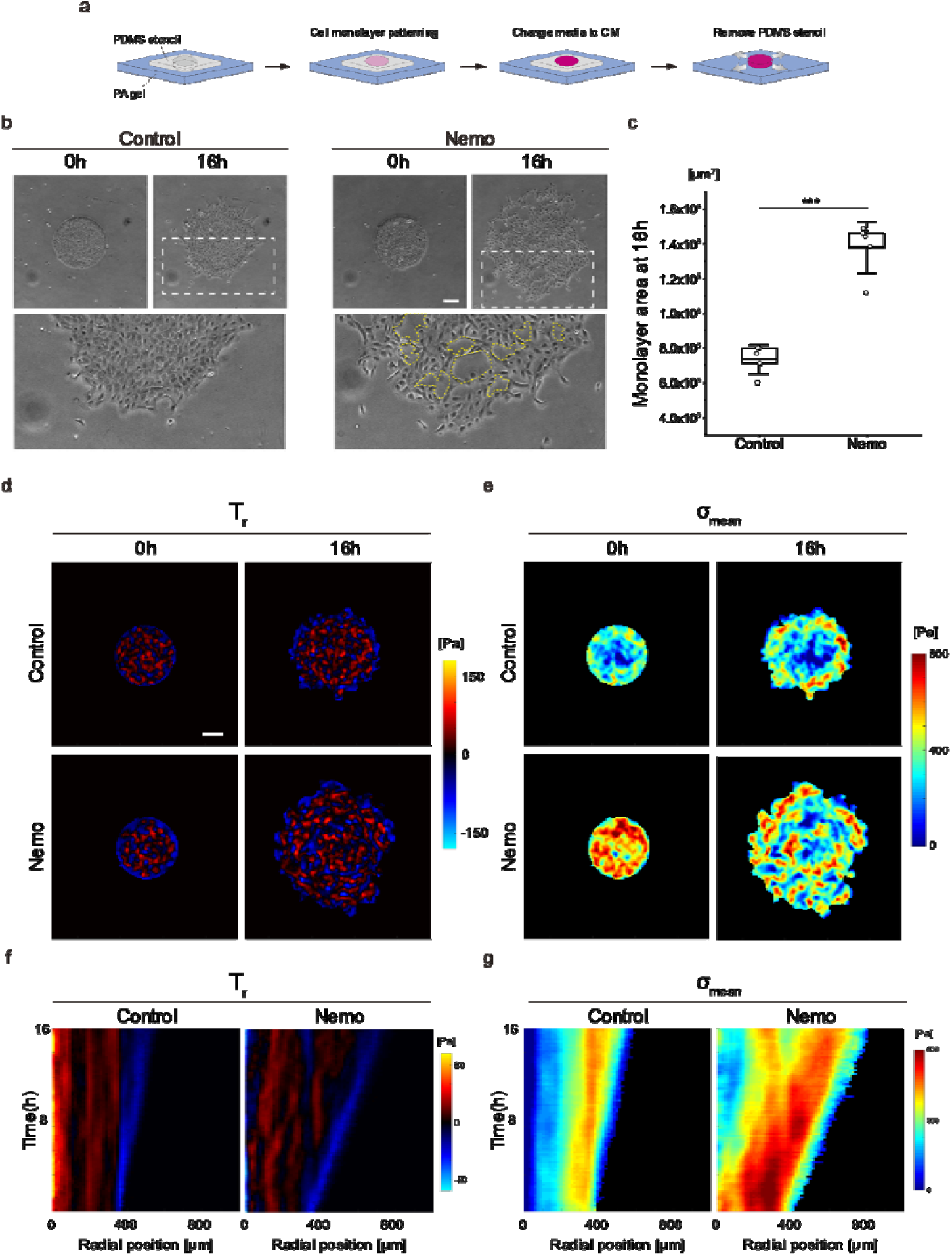
Physical analysis of normal breast epithelial cellular monolayer expansion implies the EMT of Nemo-CM-treated cells. (a) Schematics of cellular monolayer micropatterning and expansion. (b) Phase-contrast images of normal breast epithelial cellular monolayers at 0 and 16h. Yellow dotted lines represent the areas of the loss of cell-cell interactions. (c) Monolayer area at 16h. Data represent mean ± SD. n = 5 individual monolayers. (d) Color-coded maps of radial components of traction (red: outward traction, blue: inward traction). (e) Average normal intercellular stress maps. (f) Kymographs of radial traction. (g) Kymographs of average intercellular stress within the monolayer. Scale bars represent 200 µm.

To observe the temporal evolution of the traction forces, data were plotted as a kymograph (Fig. 2f). The control group showed a homogeneous pattern along over time, characterized by a dominant red-colored region, indicative of outward traction, at the core. This stable traction pattern with time indicates stable cell-substrate interactions within the monolayer. On the other hand, the Nemo-CM-treated group (Nemo group) featured heterogeneous traction patterns. Intercellular stresses were then calculated from the tractions, and the average normal stresses (σ_mean_) were plotted (Fig. 2e). Right after the free expansion was initiated (at 0 h), the control group displayed low-stress values in general and the lowest in the core of the monolayer. In contrast, the Nemo group exhibited higher stress magnitudes predominantly within the monolayer compared to the control. After 16 hours, the control group showed mostly blue spots in the center of the bulk but red spots (high tension) along the edge, which is the tension build-up by the expanding edge. However, the Nemo group showed high-tension regions at both the expanding edge and at the center of the monolayer. The observation suggests that the actively moving outward cells are predominant populations around the monolayer, inducing high tension through the cell-cell junctions. The same trend can be inferred from the average normal stress with time evolution by kymograph (Fig. 2g). The control group maintained low tension at the center, which indicates the tension by the expanding edge was not high enough to transmit to the core of the monolayer. On the other hand, the Nemo group showed high tension in most of the radial positions, and the tension near the expanding edge maintained its magnitude for a relatively long time compared to the center region. Even after 8 hours of expansion, the central tension of the Nemo group was higher than control. This observation on monolayers suggests the control and Nemo groups are significantly in different physical states by the influence of the Nemo-CM.

### Single cell-based alterations in morphology and migratory behavior support the EMT of normal breast epithelial cells via Nemo-CM

From the cellular monolayer analysis, we observed the disruption of cell-cell interactions and aggressive expansion patterns in the Nemo-CM-treated cellular monolayers, implying the cells underwent EMT. To further investigate the phenotypic transition, we quantified the morphological and migratory features of single cells from MCF10A with different CM conditions. The phase-contrast images show representative single MCF10A cells of each CM condition, and we evaluated four morphological features (Fig. 3a). The cell spreading area of each condition showed no difference. However, aspect ratio, circularity, and solidity of cells showed significant differences. These significant differences in the three shape descriptors indicate the loss of apical-basal polarity acquiring back-front polarity via Nemo-CM treatment, implying their migratory capacity. Therefore, we also quantified the migratory ability of single cells.

**Fig. 3.**
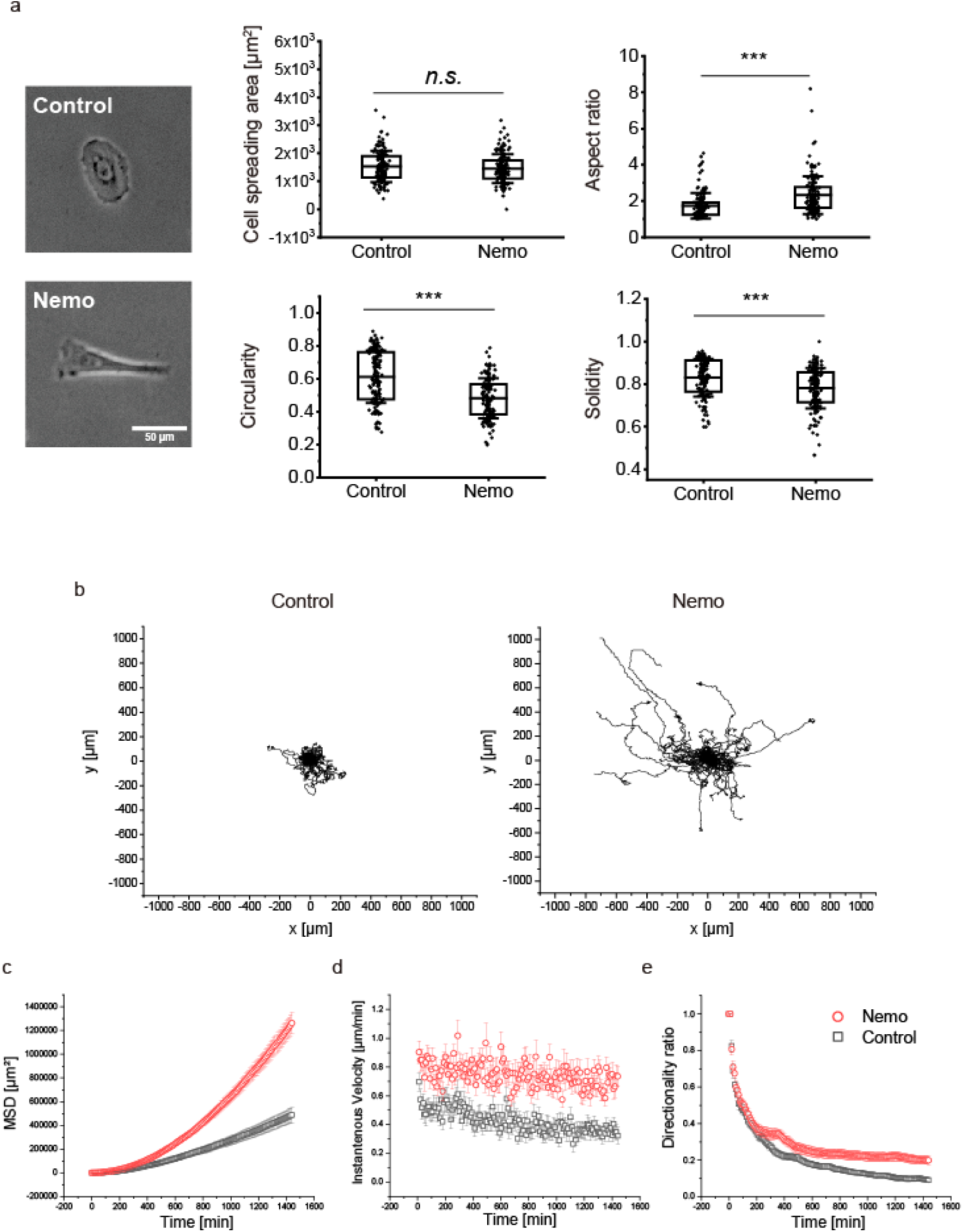
Single cell-based alterations in morphology and migratory behavior support the EMT of Nemo-CM-treated group. (a) Representative phase-contrast images of single cells under different CM conditions and morphological features. n = 139, 149 for control and Nemo, respectively, and the box plot represents mean ± SD. (b) Trajectory of single cells in each CM. (c) Mean square displacement (MSD) of single cells in each CM. (d) Instantaneous velocities of single cells in each CM. (e) Directionality ratio of single cells in each CM. n = 60 cells for each control and Nemo, from three independent experiments respectively (b-e), and data represent mean ± SEM (c-e).

The trajectories of cells under each CM were plotted, and it was obvious that single cells of the Nemo group migrated more actively than the control (Fig. 3b and Supplementary Video 2). Mean-squared displacement (MSD) was also plotted as a function of time lag. It confirmed again that the Nemo group acquired a migratory phenotype (Fig. 3c). The instantaneous velocity of single cells also supported the phenotypic transition of the Nemo group, which indicates the actively migrating characteristics of the cells (Fig. 3d). Additionally, the directionality ratio, indicating the persistence of cell migration, was also higher in the Nemo group than in control (Fig. 3e).

Collectively, the analysis of morphological and migratory features of single cells pointed out the significant phenotypic transition of cells treated with Nemo-CM. This significant shift in both morphology and motility strongly supports our speculation that the Nemo-CM is inducing an EMT in these cells.

### Biomolecular assay suggests the *SNAI1*-dependent EMT of normal breast epithelial cells in response to the Nemo-CM

To further verify whether the phenotypic transition via the Nemo-CM is the EMT, we checked the mRNA expression levels of the EMT markers (*CDH1, CDH2, VIM*) and transcription factors (*SNAI1, SNAI2, TWIST, ZEB1, ZEB2*) (Fig. 4a and Supplementary Fig. 1). Only the mRNA expression level of *SNAI1* (Snail) was significantly upregulated in the Nemo group, suggesting that the EMT via Nemo-CM is likely driven by the upregulation of the transcription factor *SNAI1* (Fig. 4a).

**Fig. 4.**
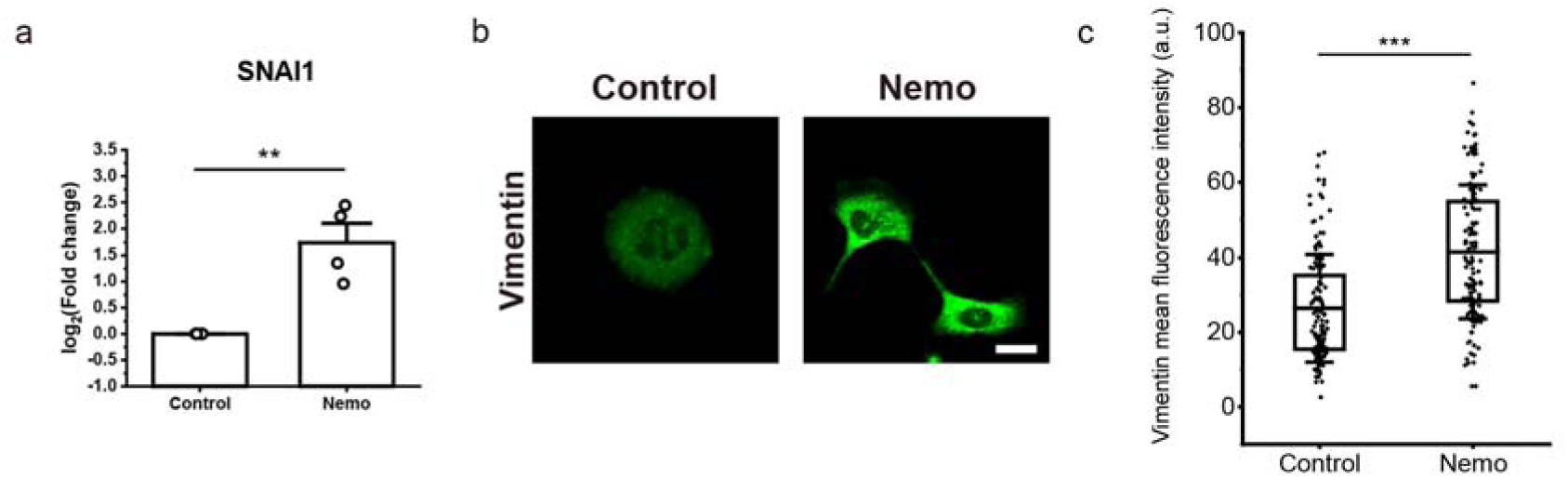
Biomolecular assay suggests the EMT of normal breast epithelial cells via Nemo-CM. (a) mRNA expression of EMT transcription factors *SNAI1*. n = 4 and data represent mean ± SEM. (b) Vimentin characterization by immunofluorescence of representative images. Scale bar: 20 µm (c) Quantification of mean intensity of vimentin. n = 141, 131 for control and Nemo respectively, and the box plot represents mean ± SD.

Although no significant differences were observed in the mRNA expression levels of *VIM* between the control and Nemo groups, we further investigated vimentin expression at the protein level using immunofluorescence in individual cells. This approach was prompted by previous findings that indicated a migratory phenotype in the Nemo group (Fig. 3b-e). Given our prior identification of the enhanced migratory ability in the Nemo group, we were able to corroborate this phenotype with immunofluorescent images as well. Regarding the mesenchymal marker vimentin, its expression was noted in both the control and Nemo groups. Indeed, when epithelial cells like MCF10A are cultured under conditions that limit their ability to form a monolayer, such as sparse culturing or any situation that restricts cell-cell contact, it can lead to an induction or enhancement of vimentin expression. However, the intensity of vimentin across cells was markedly higher in cells treated with Nemo-CM, indicating a more pronounced migratory behavior at the biomolecular level (Fig. 4b, c).

In summary, these findings collectively suggest that the Nemo-CM induces EMT primarily through the upregulation of transcription factor *SNAI1*, leading to increased vimentin expression and enhanced migratory capabilities in breast epithelial cells.

### The inhibition of EP4 suggests the PGE2-EP4-*SNAI1* axis of the EMT

The effects of PGE2 are transduced via the E series of prostaglandin receptors (EP1, EP2, EP3, and EP4). PGE2 can bind any of these four receptors, and each receptor is coupled to different signaling pathways^56^. Among these EP receptors, EP4 has received attention as a promising therapeutic target in aggressive breast cancers^56,57^. L-161,982 is a selective EP4 antagonist, and we used this inhibitor to verify whether the EMT caused by Nemo-CM is blocked. To this end, the cellular monolayers were preincubated with either L-161,982 (dissolved in DMSO, 10 μM) or the same amount of DMSO for one hour.

Even though the expansion speed of monolayers was not significantly different, the DMSO-treated controls exhibited disrupted cell-cell interactions, indicative of EMT (Fig. 5 and Supplementary Video 3). In contrast, EP4 antagonism with L-161,982 preserved these cell-cell interactions, suggesting that it effectively blocked the EMT process (Fig. 5a). These observations suggest that the elevated PGE2 in Nemo-CM initiates EMT, where this phenotypic transition may be facilitated via the PGE2-EP4-*SNAI1* axis, akin to the PGE2-EP2-*SNAI1* pathway reported by Cheng et al.^58^.

**Fig. 5.**
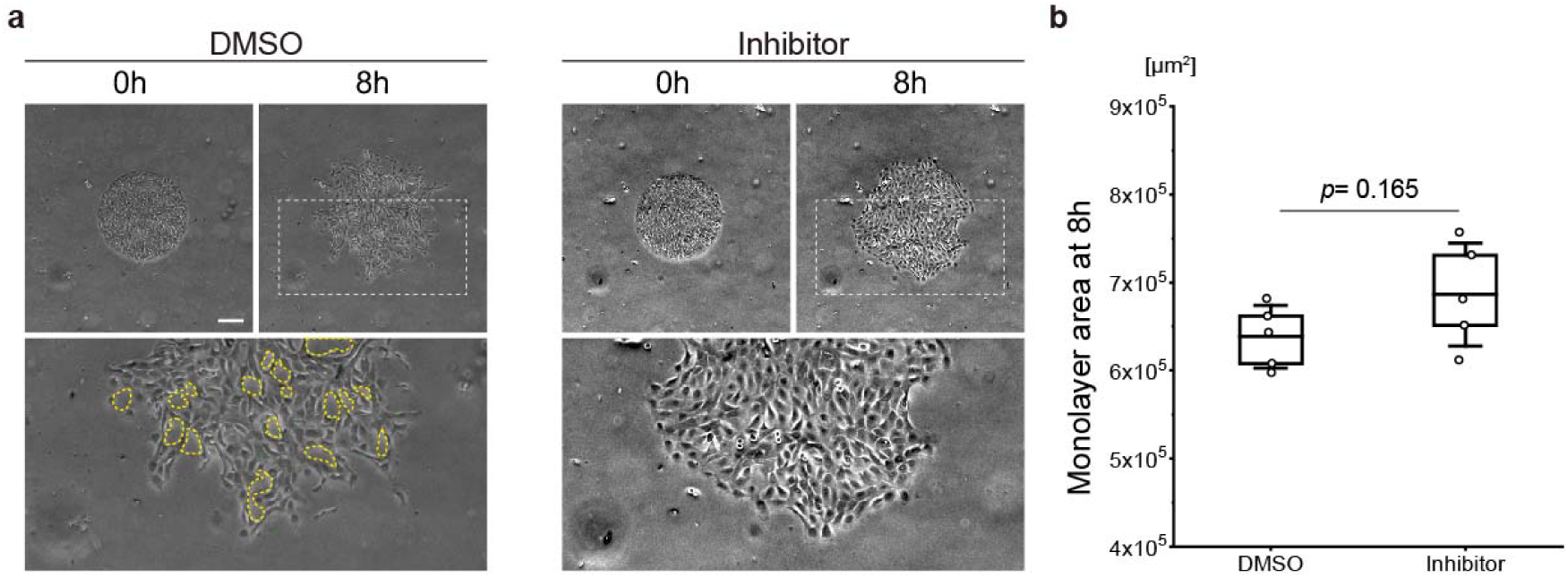
The EP4 inhibition preserves the cell-cell interactions in cellular monolayers and supports the role of the PGE2-EP4-SNAI1 signaling axis in EMT. (a) Phase-contrast images of normal breast epithelial cellular monolayers without (DMSO only) and with PGE2 inhibitor at 0 and 8 h. Yellow dotted lines represent the areas of the loss of cell-cell interactions. (b) Monolayer area at 8 h. Data represent mean ± SD. n = 5 individual monolayers. The scale bar represents 200 µm.

## Discussion

Recent findings highlighted the high *COX2* expression in the TME of various cancer types. The elevated stromal *COX2* expression have been strongly correlated with a poor prognosis in several cancers, including breast cancer. While the tumor stroma is composed of diverse cell types, there is growing evidence that *COX2* is expressed by some CAFs, known for their role in immunosuppression within tumors^25,59–62^. This suggests that *COX2*^+^ CAFs may significantly contribute to the overall *COX2* expression in the stroma. If we can decouple the *COX2*^+^ CAFs and other cells in TME, we could illuminate their unique impact on tumor progression.

Leveraging the process of nemosis to induce a state analogous to *COX2*^+^ CAFs in vitro, our study explored the paracrine influences of *COX2*^+^ CAFs on breast cancer progression. The paracrine signaling of *COX2*^+^ CAFs was simulated using conditioned media. We demonstrated that the conditioned media from *COX2*^+^ fibroblasts (the proxy of *COX2*^+^ CAFs) induced the EMT of normal breast epithelial cells, validated through both cell mechanics-based analyses and biomolecular assays. These comprehensive investigations—spanning monolayer expansion studies, quantification of single-cell biophysical attributes, qPCR for EMT markers, and vimentin immunostaining—converge to substantiate the EMT induction by Nemo-CM. Moreover, the targeted inhibition of the EP4 receptor allowed us to identify and articulate the *COX2*^+^ CAFs-PGE2-EP4-*SNAI1* axis as a central player in this process (Fig. 6), offering a plausible mechanistic link between high stromal *COX2* expression and poor cancer prognosis.

**Fig. 6.**
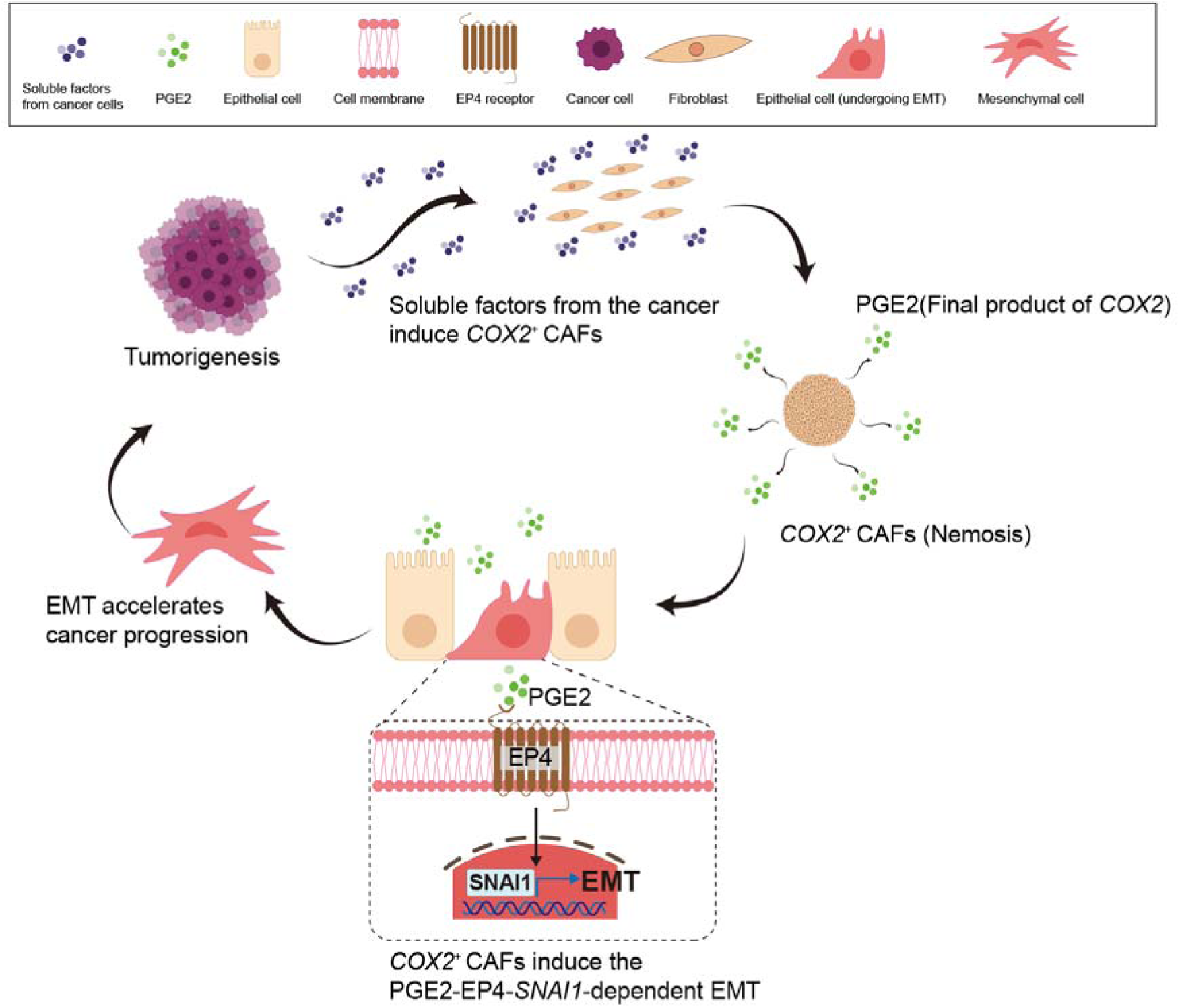
Illustration of the newly identified feedback mechanism in breast cancer mediated by *COX2*^+^ CAF via the PGE2-EP4-*SNAI1* axis.

Identifying the *COX2*^+^ CAFs-PGE2-EP4-*SNAI1* axis as a pivotal mechanism in EMT provides a novel dimension to our comprehension of TME dynamics in breast cancer. Specifically, the upregulation of *SNAI1*, one of the key regulators of EMT, underlines the critical role of transcriptional control in cancer progression. This insight does not merely echo the existing literature but also delineates the specific pathway via which *COX2*^+^ CAFs exert influence, hinting at novel therapeutic targets aimed at disrupting this pathway to counteract EMT in breast cancer.

While our in vitro study offers detailed insights into the specific role of *COX2*^+^ CAFs in the tumor microenvironment, it inherently carries some limitations. The controlled conditions of in vitro experiments allow us to decouple the complex interactions within the tumor microenvironment, providing a clear view of how *COX2*^+^ CAFs influence neighboring cells. This isolation is crucial for understanding the specific pathways and interactions at play, such as the *COX2*^+^ CAFs-PGE2-EP4-*SNAI1* axis identified in our study. However, these advantages also introduce constraints. The tumor microenvironment in vivo involves a dynamic interplay of multiple cell types, extracellular matrix components, and a host of soluble factors that cannot be fully replicated in vitro. Our model, focused primarily on the interactions between *COX2*^+^ fibroblasts and epithelial cells, does not incorporate other cellular components, notably immune cells, which play a significant role in cancer progression and response to therapy. Additionally, the use of conditioned media may not entirely mimic the direct cell-cell and cell-matrix interactions present in vivo.

To address these limitations, future research should extend these findings through in vivo studies or sophisticated 3D culture systems that better emulate the actual tumor stroma. Such studies will help validate the translational relevance of the *COX2*^+^ CAFs-PGE2-EP4-*SNAI1* axis in clinical settings and potentially guide the development of targeted therapies. Further investigations involving the interplay between *COX2+* CAFs and immune cells could also enrich our understanding of the tumor microenvironment’s complexity.

## Availability of data and materials

The datasets used and analyzed during the current study are available from the corresponding author upon reasonable request.

## Ethics declarations

### Ethics approval and consent to participate

N.A.

### Consent for publication

N.A.

### Author contributions

M.K. and S.D. conceived the research and designed experiments. M.K., S.D., and T.Y.K. performed the experiments. M.K., S.D., and J.S. interpreted data. M.K. wrote the manuscript. All authors edited the manuscript.

