## Supplemental Figure 1 for "EMT Induction In Normal Breast Epithelial Cells By *COX2*-Expressing Fibroblasts"

### Slide 1
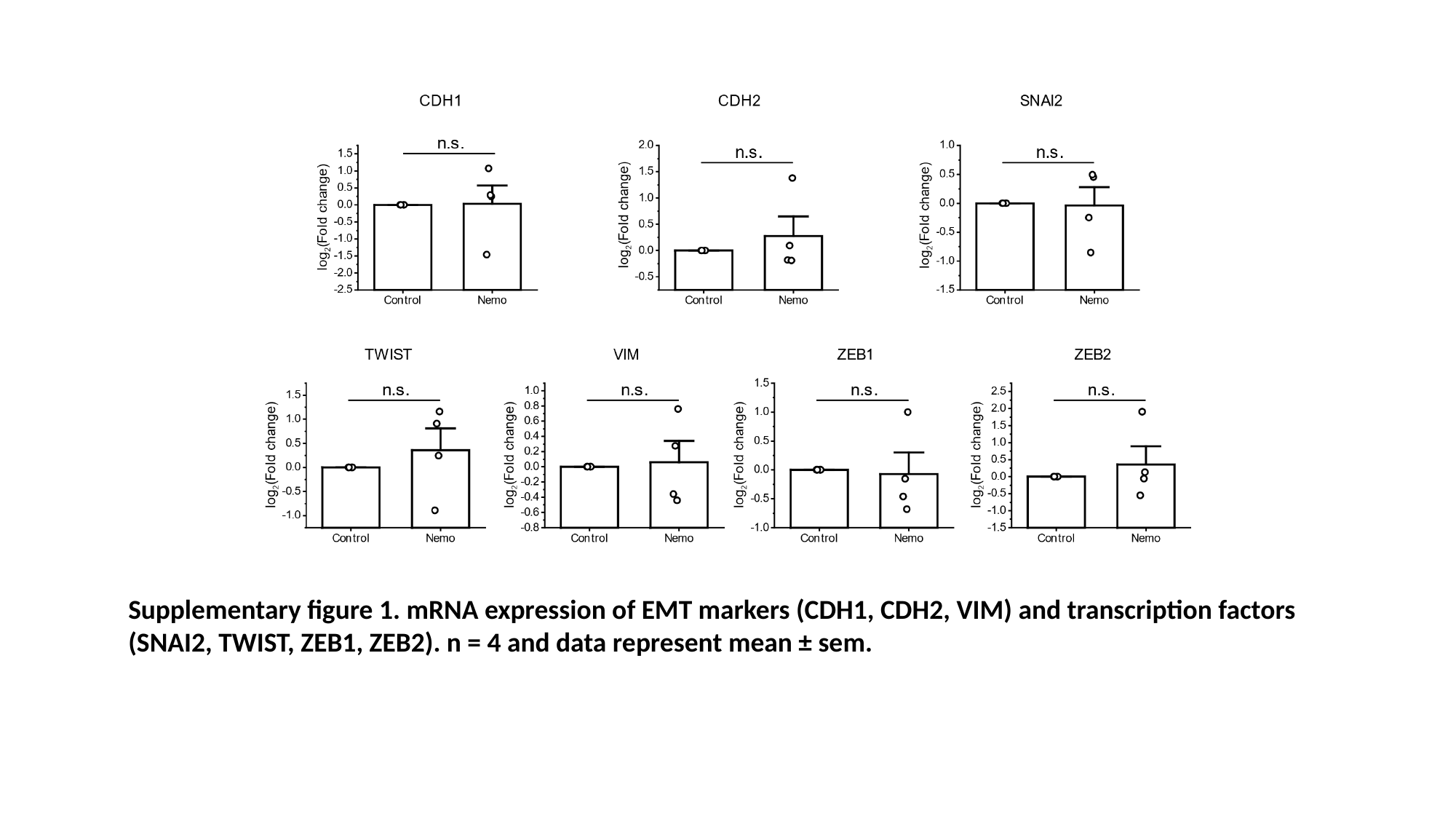

Supplementary figure 1. mRNA expression of EMT markers (CDH1, CDH2, VIM) and transcription factors (SNAI2, TWIST, ZEB1, ZEB2). n = 4 and data represent mean ± sem.
